# The Effect of Retro-inverse D-Amino Acid Aβ-peptides on Aβ-Fibril Formation

**DOI:** 10.1101/457325

**Authors:** Wenhui Xi, Ulrich H. E. Hansmann

## Abstract

Peptides build from D-amino acids resist enzymatic degradation. The resulting extended time of biological activity makes them prime candidates for the development of pharmaceuticals. Of special interest are D-retro inverso (DRI)peptides where a reversed sequence of D-amino acids leads to molecules with almost the same structure, stability and bioactivity as the parent L-peptides but increased resistance to proteolytic degradation. Here, we study the effect of DRI-Aβ40 and DRI-Aβ42 peptides on fibril formation. Using molecular dynamics simulations, we compare the stability of typical amyloid fibril models with such where the L-peptides are replaced by DRI-Aβ40 and DRI-Aβ42 peptides. We then explore the likelihood for cross fibrilization of Aβ L-and DRI-peptides by investigating how presence of DRI peptides alters elongation and stability of L-Aβ-fibrils. Our data suggest that full-length DRI-peptides may enhance the fibril formation and decrease the ratio of soluble toxic Aβ oligomers, pointing out a potential for D-amino-acid-based drug design targeting Alzheimer’s disease.

## 1. Introduction

While amino acids are, with the exception of glycine, chiral molecules, almost exclusively only the L-enantiomers are found in naturally occurring proteins and encoded in the genome. In the few cases of D-amino acids and D-amino acid-containing compounds that are seen in nature, for example, the neurotransmitter D-serine, the D-enantiomers are synthesized by enzymes and/or added as a post-translational modification.

However, cell-permeable peptides made of D-amino acids are emerging peptidomimetics with promising pharmaceutical applications. The reason for this is the resistance of peptides composed of D-amino acids to enzymatic degradation, i.e., when used as pharmaceuticals these peptides are effective for a longer time. Of special importance are D-retro-inverso (DRI) peptides which use that D-amino acids are mirror images of L-amino acids.^1^ Hence, a peptide assembled in reversed order from D-amino acids will have almost the same structure, stability and bioactivity than the parent peptide made of L-amino acids, but it will be resistant to proteolytic degradation. This combination makes DRI peptides interesting drug candidates. For instance, in one study a synthetic DRI peptide had not only structural similarity to the natural L-peptide, but it also induced a strong antibody response and had a higher resistance to trypsin than the L-peptide analog.^2^ In another recent study, Barr and coworkers showed that a DRI peptide, which mimics a 46 amino acid segment of the p53-binding domain of FOXO4, results in the release of p53 from FOXO4 and also induces cell-intrinsic apoptosis in senescent cells.^3^

In the present paper we explore the potential role of D-retro-inverso (DRI) peptides, specifically DRI-Aβ_40_ and DRI-Aβ_42_, as drug candidates targeting amyloid diseases. Marker for the neurodegenerative Alzheimer's disease are amyloid deposits in brains of patients with the disease, however, the main toxic agent may be not the final (and no longer soluble) fibrils but transient, polymorphic and soluble oligomers that could be either on-pathway or off-pathway to fibril formation. Potential drug candidates therefore should target these toxic oligomers, by either inhibiting their formation or otherwise decrease their concentration. Assuming D-retro-inverso Aβ_40_ (DRI-Aβ_40_) peptides and D-retro-inverso Aβ_42_ (DRI-Aβ_42_) peptides to form similar assemblies as (L-) Aβ_40_ and (L-) Aβ_42_ peptides, respectively, one can conjecture two mechanisms by that the DRI peptides could reduce the concentration of toxic Aβ-oligomers. First, built into the oligomers they may induce an antibody response cleaning away the oligomers. Another possibility would be a higher stability and resistance to proteolytic degradation of hybrid fibrils, shifting the equilibrium away from the toxic oligomers toward the less toxic fibrils. Both mechanisms require them to form hybrid aggregates with L-Aβ-peptides. The purpose of this paper is to evaluate whether such hybrid fibrils can form and if they are stable.

The existing applications of D-retro-inverso proteins as inhibitors are possible because these molecules share the geometry and stability of the L-parent. However, the structures of the two kinds of proteins are not identical. For instance, a helix will have in a DRI-protein the opposite winding than in the original L-protein. These dissimilarities may lead to subtle differences in structure and stability of fibrils built from retro-inversed D-Aβ peptides. For this reason, we start our investigation by probing in all-atom molecular dynamics simulations the stability of two Aβ_40_ and Aβ_42_ fibril models deposited in the Protein Data Bank (PDB), and we compare their stability with that of the corresponding DRI versions. We find that the DRI forms may vary in the twist of β-sheets; and in some cases, they have lower stability due to interaction of side chains with end groups. In the second part we then test the effect of DRI-Aβ peptides on fibril formation. We observe that hybrid assemblies of DRI-peptides with L-peptides have comparable stability with that of L-fibrils. Hence, there is a likelihood for the cross fibrilization of L‐ and DRI‐Aβ peptides, which implies that full-length DRI-peptides may enhance the fibril formation and decrease the ratio of soluble toxic Aβ oligomers.

Previous studies focused on inhibitors formed from short peptides containing D-amino acids,^4.5-7^ which decrease formation of fibrils^8^ or even dissolve them.^6^ The exception is a recent study of fibril formation of full-length D-enantiomer of Aβ_42_ (with nonreversed sequence), which showed that the mixing of L‐ and D-peptides enhances fibril formation^9^ and reduces the concentration of soluble toxic Aβ oligomers, protecting PC12 cells. The common theme in all these previous studies is that replacing a single L‐residue by a D-amino acid already alters significantly the binding affinity.^10^ For example, the Daggett group has designed short peptides with alternating D‐ and L-amino acids that bind to toxic Aβ_42_-oligomers and reduce their toxicity.^11,12^ The D-peptides D(pgklvya) and D(kklvffa) which are based on the segment made of residues 16-20 in Aβ peptides, inhibit fibrillogenesis of Aβ_42_ and increase the lifetime of transgenic *Caenorhabditis elegans* model.^13^ These peptides act also in their L-form as inhibitors (i.e. share the binding affinity) but have the additional advantage of increased protease resistance that motivates also our work. A DRI‐ Aβ-fragment, Ac-D(rgffvlkgr)-NH2 named RI-OR2, was used as an inhibitor of Aβ oligomerization,^14-17^ and shown to be more effective than the L-form.

Unlike this previous work, we study in this paper aggregates of the full-length DRI-Aβ peptides. Our results give not only a detailed comparison of L‐ and DRI-fibrils at atom level, but also suggest a role for DRI-Aβ based drug design targeting Alzheimer’s disease.

## 2. Methods

### 2.1 Model construction

In this study, we compare structure and stability of two experimentally resolved Aβ-fibril models with that of their DRI analogs, and with that of hybrid fibril models made from a mixture of DRI‐ and L-peptides. The first model is a two-layer U-shaped Aβ_40_ model (PDB ID: 2LMN),^18^ while we choose a recently resolved two-layer S-shape fibril model (PDB ID:2NAO) for Aβ_42_.^19^ The two models were not only chosen because we are familiar with them from our previous work^20^ but also because they allow us to test whether changes in stability and structure resulting from replacing L-peptides with DRI peptides differ for U-shaped and S-shaped motifs. Representative conformation of the two-layer Aβ_40_ and Aβ_42_ fibrils are shown in Figure 1. The first eight residues in the Aβ_40_ model have not been resolved in the PDB structure, they are added here assuming these residues to be in a random configuration. In previous work, we have shown that the minimal stable fibril fragment of S-shaped Aβ_42_ chains is a hexamer,^20^ suggesting a minimum number of six chains per layer for our Aβ_42_ model. For consistency, we chose the same number of chains per layer for the Aβ_40_ model. However, for the hybrid fibril models, built by mixing L-peptides with DRI peptides, we prefer an odd number of chains per layer, and here each layer consist of seven chains, allowing combinations such as 3xL-1xDRI-3xL. Finally, in order to see whether the ability to form two-layer fibrils differ between the original L-fibrils and the DRI-fibril (or hybrid fibril) models, we also generated corresponding one-layer Aβ_42_ fibril models (one-layer Aβ_40_ fibrils have not been observed and were therefore not considered) by removing the residues of one layer. In order to study the differences in the interaction of side chains with the end groups in L and DRI fibrils, we also considered mutants K28A and K28E, and such with capped end groups.

**Figure 1.**
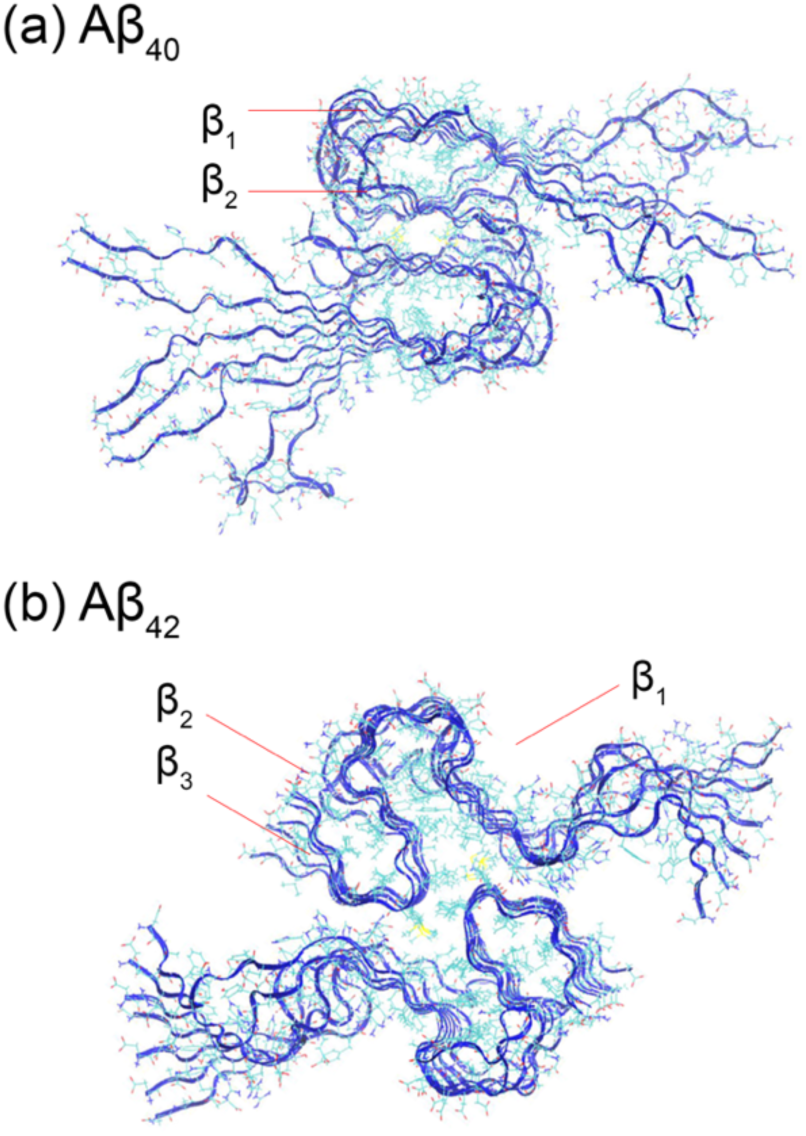
Two-layer Aβ_40_-fibril (a) derived from the experimental model with PDB-ID 2LMN, and (b) the corresponding model of Aβ_42_ fibrils as derived from the PDB-structure 2NAO. The individual chains in the two models are L-peptides.

The retro-inverso D-peptides (DRI-peptide) differ from their L-peptide parents in that they have a reversed sequence *and* L-amino acids replaced by D amino acids. As the position of the carbon and oxygen atoms are with respect to the peptide bond nearly symmetric to the position of the nitrogen and hydrogen atoms (with only a small difference between the C-O and N-H bond length and the bond angle), see Figure 2, an initial DRI fibril structure is easily derived from the L-parent by replacing in the PDB-file the names of backbone atoms as follows: N->C, C->N, H->O, O->HN (HN represent a hydrogen atom that is bonded with a N atom). This substitution changes the direction of the chain; and as the positions of side chains are not altered, turns L amino acids into D amino acids. The N-HN and C-O bond length and bond angles in the resulting configuration are incorrect, but they assume correct values after minimization and a short NPT simulation.

**Figure 2.**
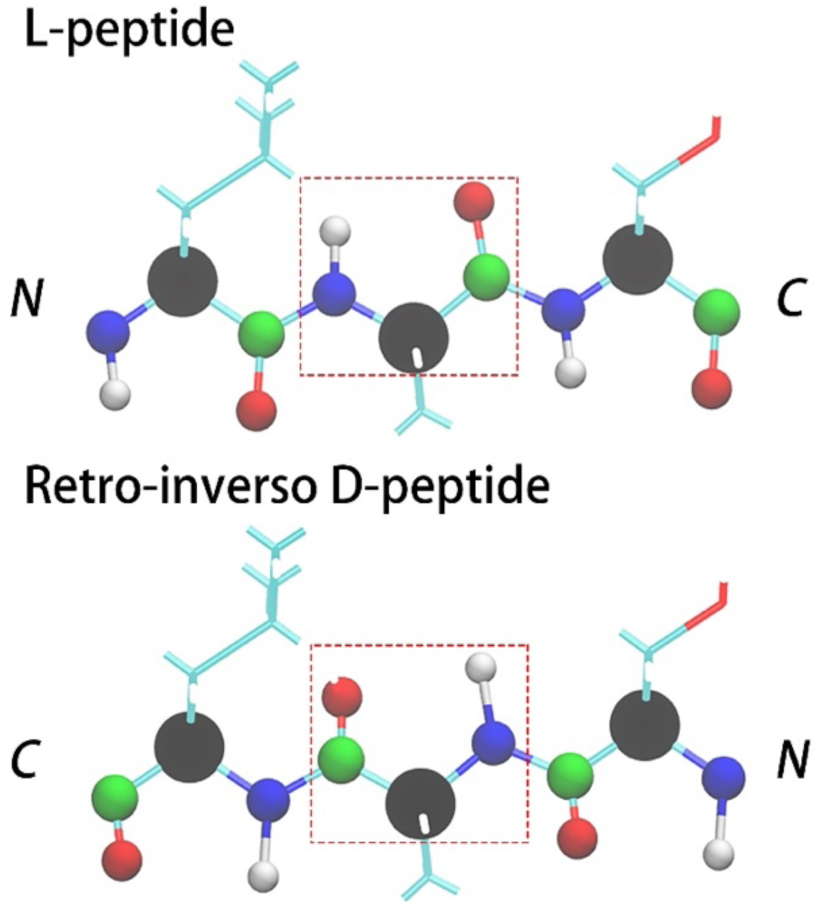
Backbone replacements that converts an L-peptide into its retro-inverso DRI form made of D-amino acids. The backbone atoms are shown as spheres using the color code: CA-black, N-blue, HN-white, C-green and O-red.

L-peptide fibrils and DRI fibrils may differ in the twist of their β-strands, and there is a danger that the above procedure “freezes” into configurations with the wrong twist. In order to ensure that our procedure does not lead to hidden biases, we also back-generated, using AmberTools,^21^ L-fibrils from our DRI-fibrils, and compared their stability and structural changes with the originals. Hybrid fibrils, containing both L-peptides and their DRI analogs, are constructed by using the above procedure only for selected chains.

Representative conformations of hybrid models made of L and DRI Aβ_40_ peptides are shown in supplemental Figure S1 (a,b,c) where the L-peptides are colored in blue and DRI-peptide in red. A list of all considered models is shown in supplemental Table S1. In order to compare more easily the DRI-peptides with the L-parent we number the residues in DRI peptides starting from the C-terminus, not from the N-terminus as is the usual convention. In this way, residue X is the same type in both L-peptide and DRI peptides (except either a L-amino acid or a D amino acid). Note also that in tables, figures, and throughout the text we use in fibril names the abbreviation L for L peptides, D for DRI peptides, L’ for the K28A mutant of a L-peptide, L* for the K28E mutant of a L-peptide, D’ for the K28A mutant of a DRI peptide, and D* for the K28E mutant of a DRI peptide. Hence, 7-L/7-L names a two-layer fibril where each layer is built of 7 L-peptides, while 7-D* would be a single-layer fibril made of seven DRI peptides, each with the mutation K28E.

### 2.2 Molecular dynamics simulations

Our simulations rely on the GROMACS 5.1.5^22^ program package using the CHARMM force field (version: 36, jul-2017), which includes non-standard amino acids such as the 19 D-amino acids,^23^ and TIP3P water^24^. The minimum distance between peptides and the cubic solvent box edge is set to 0.8 nm in the initial conformation, and the solvent is neutralized by Na+ and Cl−ions. All models are minimized and thermalized over 5ns runs of molecular dynamics in an NVT ensemble, followed by a run of the same length in a NPT ensemble. Because of the use of periodic boundary conditions are electrostatic interactions calculated with the particle mesh Ewald(PME) method^25^. The cutoff of van der Waals(vdW) and electrostatic interactions are 1.2/1.2 nm which is suggested for CHARMM force field 36. Keeping the bond length fixed with the LINCE^26^ and SETTLE algorithms^27^ allowed for an integration time step of 2 fs. The temperature of 310K and a pressure of 1 bar are controlled by v-rescale thermostat^28^ and Parrinello-Rahman barostat.^29^. For each system, we have performed two independent runs of 50ns, using only the last 25ns for analysis. Most of the analysis uses the tool-set provided with GROMACS. However, representative configurations are visualized with VMD^30^, a program which we also use to generate the Ramachandran plots.

## 3. Results and discussion

### 3.1 Comparison of L-peptide fibrils with DRI-peptide fibrils

AFM measurements have shown that Aβ-peptides, made of D amino acids, can form amyloid fibrils.^9^ However, while D-retro-inversed proteins fold into the same structures as the L-forms, and have comparable stabilities, there are differences, for instance, in helix-winding. Hence, it is not guaranteed that DRI-Aβ peptides assemble into the same kind of amyloids than the L-parents, and if, whether the DRI-fibrils have comparable stability. This is important to know when considering a role of DRI-Aβ as potential drugs targeting Alzheimer's disease. Hence, we start our investigations by looking first into the stability of experimentally derived fibril models, replacing the L-Aβ chains with DRI-Aβ peptides. For this purpose, we follow the time evolution of each DRI fibril model for 50 ns in two independent molecular dynamics simulations at T=310 K, and we compare the trajectories with that of corresponding L-fibril model runs. Construction of the two-layer DRI-Aβ_40_ and DRI-Aβ_42_ fibril models is described in the Method section. As fibrils with S-shaped L-Aβ_42_-peptides have been observed not only as two-layer assemblies but also as single-layer aggregates, we also looked into the stability of the single-layer DRI-Aβ_42_ fibrils.

#### 3.1.1 Double-layer Fibrils

We show in Figure 3 the root-mean-square-deviation (RMSD) as function of time for the two double-layer fibril architectures, built from either L or DRI peptides. We ignore in the RMSD calculation the flexible segment made of residues 1 to 10 in each L-Aβ_42_ chain, and the corresponding segment in the DRI peptides. In a similar way we ignore likewise the flexible residues 1 to 8 for Aβ_40_, ensuring that in both cases the RMSD calculation goes over 32 residues. Out of the two runs for each system, we plot here (and in all following figures) only the run that lead to the higher final root-mean-square-deviation. The RMSD time evolution, and visual inspection of snapshots sampled in the respective trajectories, indicate that the L-forms and the DRI-forms of both the Aβ_40_ and the Aβ_42_ two-layer models have similar stability. Representative configurations of both L and DRI Aβ_40_ fibril configurations are shown in Figure 4(a, b). In order to make the picture more easily readable only one of the two layers is displayed. The Ramachandran plots of the phi/psi angle distribution in Figure 4 (c, d) of the L and DRI form confirm that our force field provides a faithful representation of D-amino acids.

**Figure 3:**
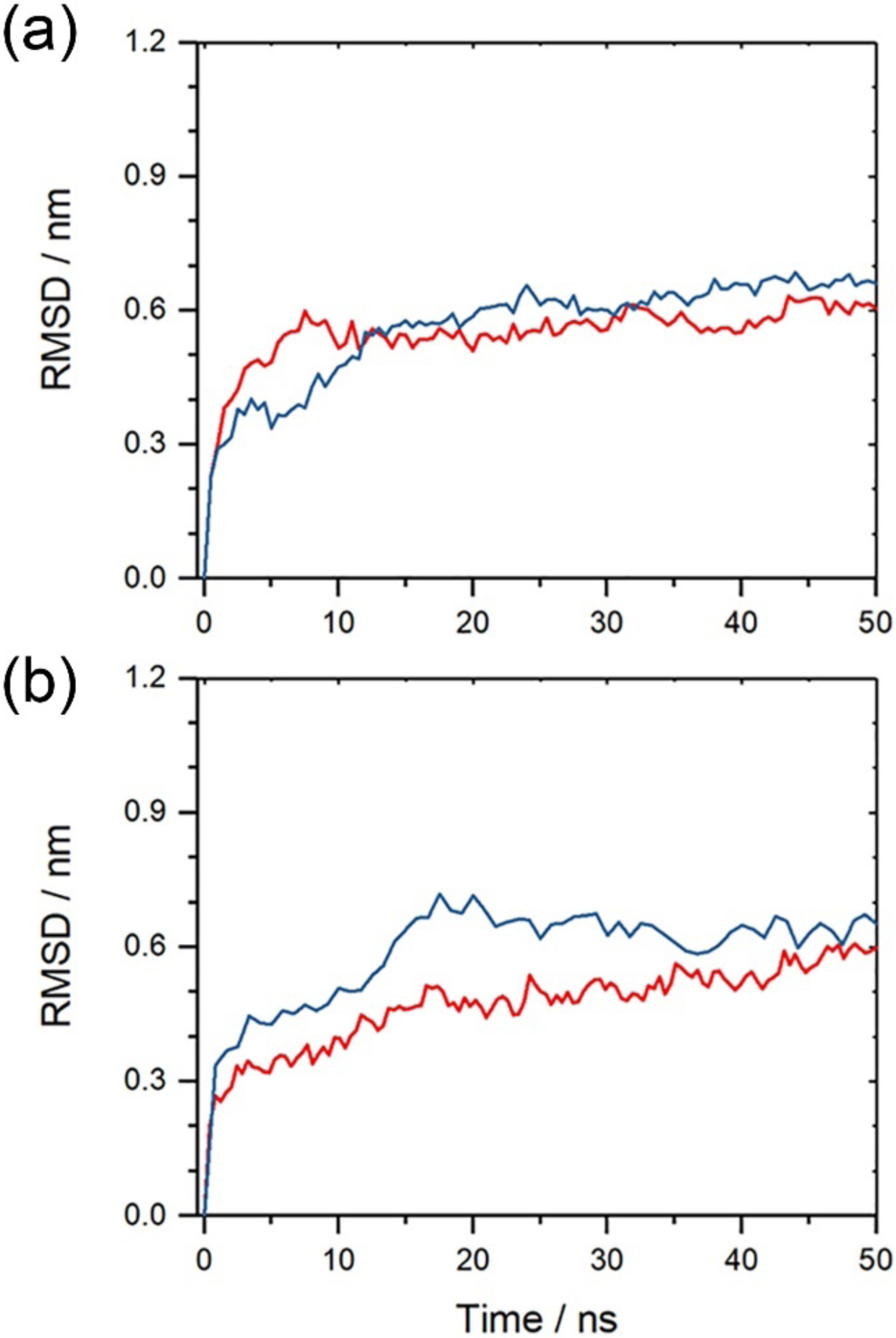
Root-mean-square deviation (RMSD) to the start configuration as function of time for (a) two-layer Aβ_40_-fibrils and (b) two-layered Aβ_42_-fibrils. Data for fibrils made from L-amino acids are drawn in red; such for fibrils built from DRI-peptides are drawn in blue. Note that the flexible first eight N-terminal residues in L-Aβ_40_, and the corresponding residues for the DRI-peptides, are not considered in the RMSD calculation. Shown is for each model the run that lead to the larger RMSD.

**Figure 4.**
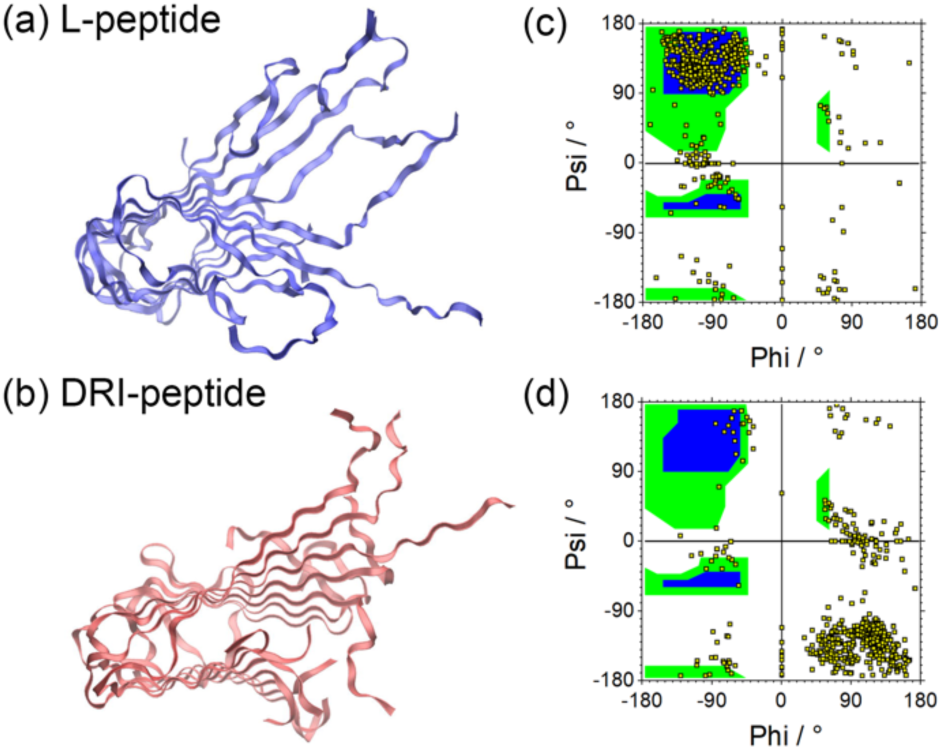
Representative conformations of a single layer of the two-fold Aβ_40_ fibrils built from in L-amino acids (a) and such built from DRI peptides(b). The corresponding Ramachandran plot are shown in (c) and (d).

The β-sheets in most fibril models are twisted along the fibril axis. This twist can be described by an angle between β-sheets in adjacent chains along the fibril axes. For Aβ_40_, this twist angle can be defined as the dihedral angles between Cα-atoms of residue 16 and 20 on neighboring chains, and for for Aβ_42_ as the dihedral angles between Cα-atoms of residue 25 and 29, located in the β2 region. Our convention is that the twist angle of L-peptides is positive. Naively, one would expect from the Ramachandran plot that the DRI-fibrils have a twist angle which is of opposite sign but equal magnitude than the one seen in the corresponding L-fibrils. However, while this appears to be the case for the Aβ_42_ fibril model, it is not the case for the Aβ_40_ fibril model, see Table 1.

**Table 1.** The twist angle between β-sheet in two-layer Aβ_40_ and Aβ_42_ models. The twist angles are defined as dihedrals of Ca atoms of residue 16 and 20 for Aβ_40_ and residue 25 and 29 which are located on β2 region for Aβ_42_. Shown are both the values for the first-layer and the second-layer (separated by “/”); the averages and standard deviations are taken over both trajectories.

| Double-layer $A\beta_{40}$ | | | | | | | | |
| --- | --- | --- | --- | --- | --- | --- | --- | --- |
|  | L-twist initial conformation |  |  |  | D-twist initial conformation |  |  |  |
|  | 1<sup>st</sup> run | 2<sup>nd</sup> run | Average | STD | 1<sup>st</sup> run | 2<sup>nd</sup> run | Average | STD |
| 7-L/7-L | 13.7/12.5 | 10.4/12.4 | 12.2 | 1.2 | 9.9/10.6 | 11.4/8.9 | 10.2 | 0.9 |
| 7-D/7-D | -4.5/-4.9 | -9.4/-3.1 | -5.5 | 2.4 | -4.4/-5.2 | -5.5/-5.1 | -5.1 | 0.4 |
| Double-layer $A\beta_{42}$ | | | | | | | | |
| 7-L/7-L | 1.2/2.1 | 2.7/1.6 | 1.9 | 0.6 |  |  |  |  |
| 7-D/7-D | -2.7/-0.9 | 1.9/-0.1 | -0.4 | 1.6 |  |  |  |  |

Compared to the L-peptides (12.2 degree), the DRI-peptides (−5.5 degree) have twist angles of much smaller magnitude. This difference is not an artifact of the way our DRI fibrils are constructed which leads to an initial twist angle of *same sign* and magnitude than seen in the L-fibril. However, enforcing in the initial fibril configuration a twist angle that has the same magnitude than the one in the L-fibril, but the correct *opposite sign,* does not change the final twist angles at the end of the trajectories. In a similar fashion does the final twist angle of a L-fibril not change if the initial twist angle is set to is that of the DRI-fibril. Hence, L-Aβ_40_-fibrils and DRI-Aβ_40_-fibrils have not only, as expected, twists of opposite sign, but they also differ in the magnitude of twist.

This effect may not be observed for the Aβ_42_-fibrils because the S-shaped chains form three strands that are shorter than the two strands in the U-shaped chains in Aβ_40_-fibrils. Because of the shorter length the twist angle between neighboring chains is not well-defined for the Aβ_42_-fibrils, and it is therefore difficult to distinguish the magnitude of twist angles between L and DRI fibrils. While for the two-layer Aβ_40_ fibrils the twist changes from around 12° for L-peptides to about −5° for DRI peptides, the change is from about 2° (1°) (L-peptides) to −0.5°(2°) (DRI peptides) for the two-layer Aβ_42_-fibril (i.e., compatible with a twist angle of zero degree).

An interesting observation is that the number of inter-chain hydrogen bonds (listed in supplemental Table S2) differs between L and DRI fibrils. When calculating the number of inter-chain hydrogen bonds, we ignore again for the Aβ_40_ fibril architecture the flexible residues 1-8. We find for the double-layered L-fibrils on average 32(4) hydrogen bonds between the U-shaped chains, 16 on each side. However, the four chains at the end of the fibril fragment are connected by only 13(3) hydrogen bonds with their neighbors. On the other hand, if the fibril is made of DRI peptides, the corresponding numbers are with 25(4) hydrogen bonds between chains within the fibril (12-13 on each site) lower than the ones found for the L-fibril. The missing ≈7 hydrogen bonds are mostly at the end of the β1-strand (in the region around residues 23-29). The decrease in the number of hydrogen bonds is larger for the end chains, which on average are connected by only 8(3) hydrogen bonds with their neighbors. The lower number of hydrogen bonds in DRI fibrils than seen in the L-parent fibrils may explain the small differences in stability observed between the two fibrils, with the larger loss for the end chains suggesting a longer nucleation phase for DRI peptides than seen for L-peptides.

The loss of hydrogen bonds is smaller for the double-layer Aβ_42_ fibril architecture with its S-shaped chains. Here, we measure only interchain hydrogen bonds involving residues 11-42, i.e., ignoring again the flexible residues 1-10. In these fibrils are chains connected by 43(3) hydrogen bonds, about 22 on each site, if the fibril is made out of L-peptides, and by 38 (4), about 19 on each site, if the fibril is assembled from DRI peptides. The numbers are again lower for the four end chains: 20(3) for L-fibrils and 15(2) for DRI fibrils. The missing ≈5 hydrogen bonds are again located at the end of the β1-strand, around residues 19-24. As in the case of the two-layer Aβ_40_ fibrils is the reduction in the number of inter-chain hydrogen bonds correlated with a slight loss of stability of the DRI-fibril, but the smaller loss of hydrogen bonds in the two-layer Aβ_42_ fibril architecture does not translate into a smaller loss of stability for the DRI fibril.

It is tempting to connect the disparity in the number of inter-chain hydrogen bonds between L and DRI fibrils with the earlier noticed difference in the magnitude of the twist angle, which is also larger for Aβ_40_ than for Aβ_42_. The correlation between interchain hydrogen bond number and twist angle suggests that the twist angle is connected with the different geometries and symmetries of the U-shaped Aβ_40_ and S-shaped Aβ_42_ fibril models.

In the U-shaped Aβ_40_ fibrils the β1 and β2 strands are staggered, and a for the U-shape geometry characteristic salt bridge D23-K28 connects chain (i) and (i+2) in L-fibrils, but chains (i) and (i-2) in DRI fibrils, see also Figure 8e in a later section of this paper. This salt bridge is stable in our trajectories, and it constraints the geometry and movements of the chains in the fibril. In the L-fibril this salt bridge does not interfere with the intrinsic twist that is seen in a β-strand (about 10°) nor does it restrict the formation of hydrogen bonds involving residues 23 to 29. On the other hand, as in DRI peptides the salt bridge is between chains (i) and (i-2), it interferes with formation of hydrogen bonds between chains (i) and (i+1) involving residues 23-29, in this way hindering the emergence of the “natural” twist in the β-strands.

The problem does not exist for the S-shaped Aβ_42_ fibrils where the chains align side-by-side allowing for a well-defined intra-chain salt bridge to from between residues K28 and A42. Not only are the three β-strands too small to develop a pronounced twist, there is also less of a difference in the constraining effect by the inter-chain salt bridge. When two such S-shape fibrils pack into 2-layer models, the inter-chain interactions on the packing surface enhance further the stability of fibrils and decrease its flexibility reducing the twist even more. A switch from L-peptides to DRI peptides will not change this scenario for the S-shaped Aβ_42_ fibrils, however, it will still lead to a smaller number of hydrogen bonds in the DRI-fibril than seen in the L-fibril.

The above noted numbers of inter-chain hydrogen bonds, and the root-mean-square-deviations in Figure 3, indicate that for the two-layer Aβ_42_ fibril architecture the stability of L fibrils and DRI fibrils is similar, with only slightly higher RMSD values measured for the DRI fibrils. Note, however, that in the two-layer model, the interaction between two-layers increases the stability of β2-region and that of the turn, which in turn restraints the movement of β3. As a consequence, the stability of the two-layer Aβ_42_ fibril may be so large that differences in stability between L and DRI form are difficult to notice.

#### 3.1.2 Single-layer Aβ_42_ Fibrils

This hypothesis is supported by the number of interchain hydrogen bonds measured for the single-layer fibril. On average, we observe for L-fibrils 42(4) interchain hydrogen bonds, about 21 on each site, with a slightly lower number of 20(3) hydrogen bonds connecting chains at the end of the fibril. Both values that are comparable to the numbers seen for the two-layer fibrils. On the other hand, for the single-layer DRI fibrils we find only 35(4) hydrogen bonds, 13 on each site, and 15(2) for the end chains, much lower numbers than the values seen for the two-layer system. As for the double-layer fibril are the missing hydrogen bonds located at the end of the β1-strand, around residues 19-24.

These much lower numbers of inter-chain hydrogen bonds in the DRI fibril than seen in the L-fibril indicate that the single-layer DRI fibrils have a much lower stability than the corresponding L-fibrils. This is supported by Figure 5 which shows the time evolution of the root-mean-square deviation measured in simulations of single-layer Aβ_42_ fibrils. Here, significant differences between L and DRI fibrils are seen, leading to noticeable larger final RMSD values for the DRI-fibril. Visual inspection of the structures shown in Figure 6a and 6c indicates that while the overall fibril structure is conserved, and inter-chain hydrogen bonds have not been broken, the β3-region does not pack as well with the β2-section in DRI-peptides as they do in L-peptides fibrils. This is an important observation as the β2-turn-β3 region is the main hydrophobic region and critical for the stability of the fibril.^20^ The twist angles differ more between L and DRI fibrils for single layer Aβ_42_-fibrils than they do for the double layer fibrils. However, with twist angles of about 5° for L peptides versus −10° for DRI peptides is the difference still small, see Table 2.

**Table 2.**
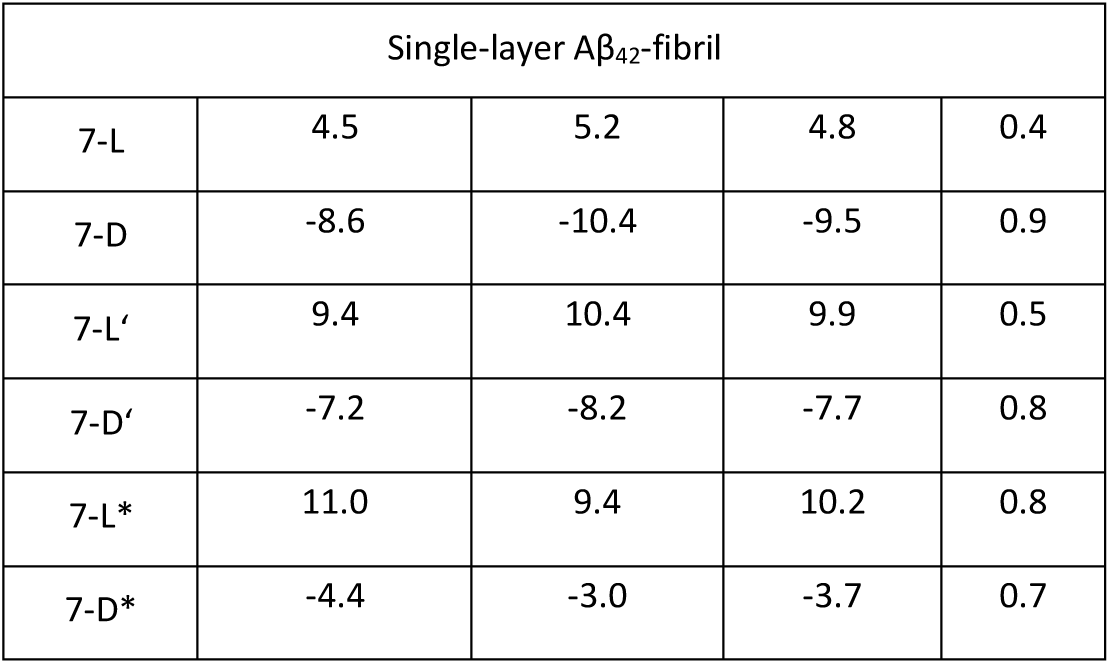
The twist angle between β-sheet in single-layer Λβ_42_ models. The twist angles are defined as dihedrals of Ca atoms of residue 25 and 29 which are located on β2 region for Λβ_42_.

| Single-layer $A\beta_{42}$-fibril | | | | |
| --- | --- | --- | --- | --- |
| 7-L | 4.5 | 5.2 | 4.8 | 0.4 |
| 7-D | -8.6 | -10.4 | -9.5 | 0.9 |
| 7-L' | 9.4 | 10.4 | 9.9 | 0.5 |
| 7-D' | -7.2 | -8.2 | -7.7 | 0.8 |
| 7-L* | 11.0 | 9.4 | 10.2 | 0.8 |
| 7-D* | -4.4 | -3.0 | -3.7 | 0.7 |

**Figure 5:**
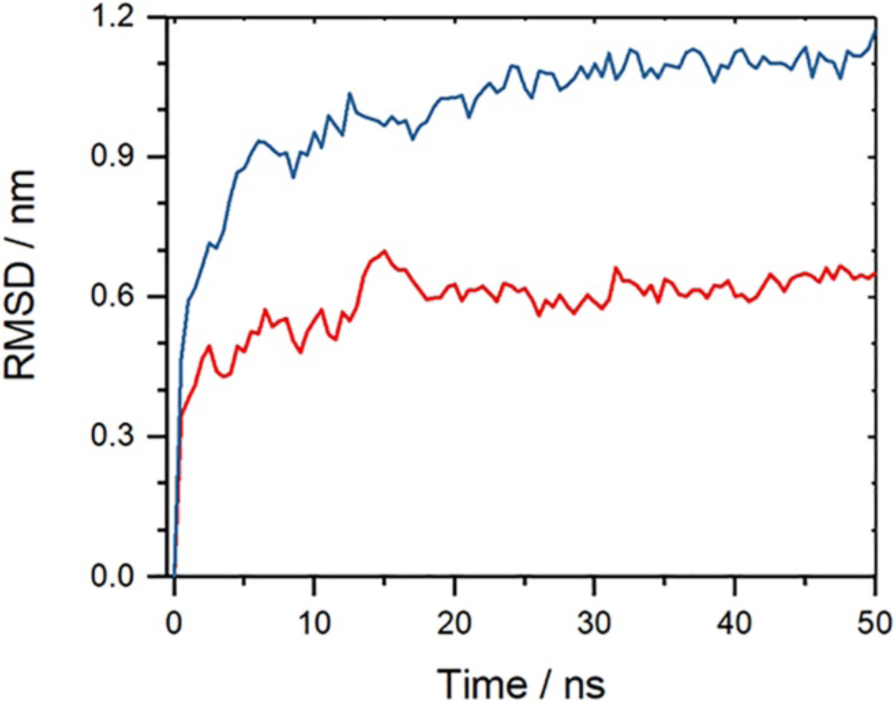
Root-mean-square deviation (RMSD) to the start configuration as function of time for single-layer Aβ_42_-fibrils. Shown is for each model the run that lead to the larger RMSD. Data for fibrils made from L-amino acids are drawn in red; such for fibrils built from DRI-peptides are drawn in blue. The flexible first ten N-terminal residues in L-Aβ_42_, and the corresponding residues for the DRI-peptides, are not considered in the RMSD calculation.

**Figure 6:**
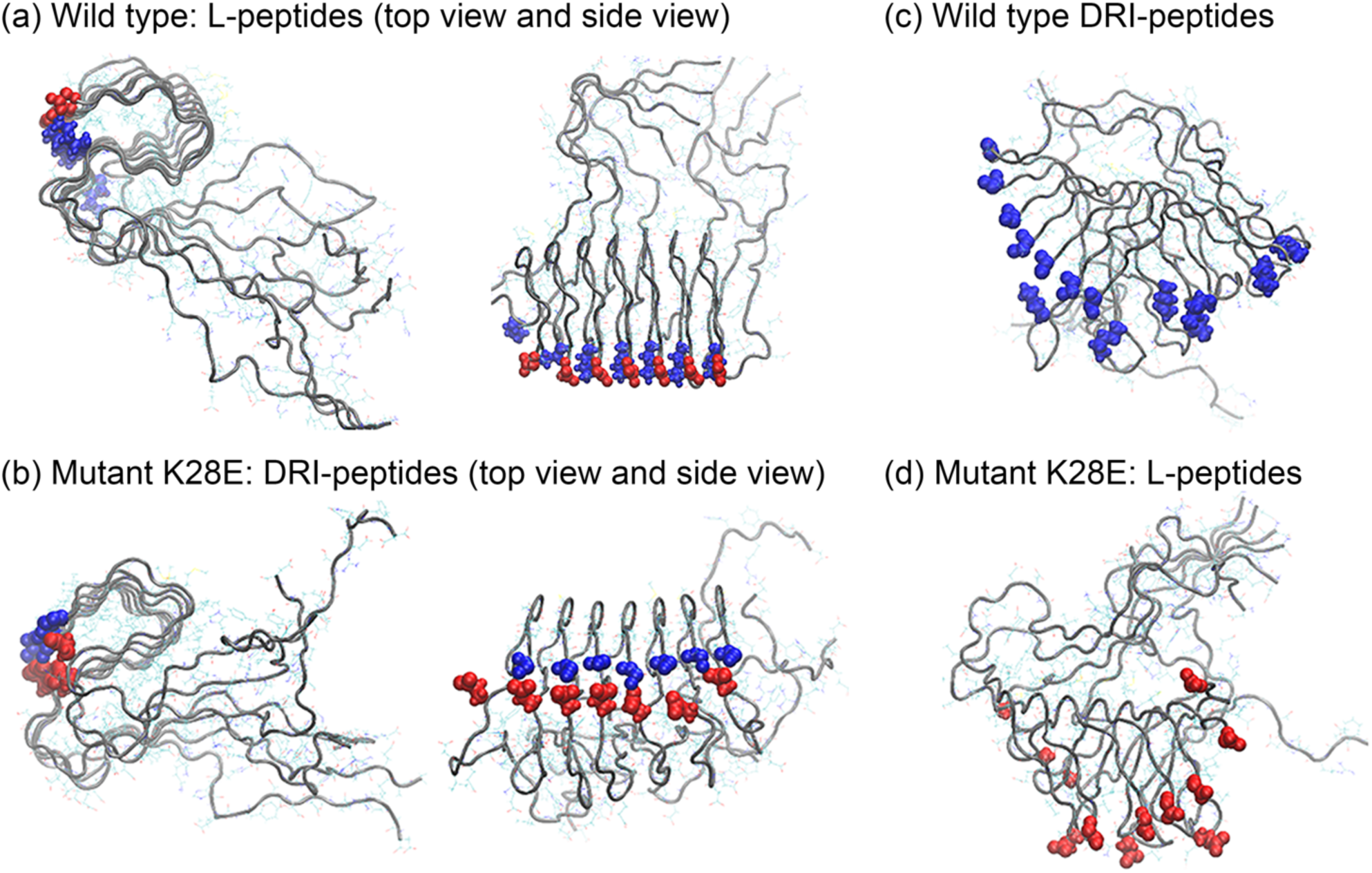
Representative conformation of single-layer-fibrils made of wild type (a) and mutant K28E (b) L-Aβ_42_ chains. The corresponding conformations for DRI-Aβ_42_-fibrils are shown in (c) for the wild type and in (d) for the K28E mutant. Blue spheres mark presence of a positive charge and red ones of a negative charge.

The disparity in structure and stability between single-layer L and DRI Aβ_42_ fibrils may be caused by the intra-chain salt bridge that in the L-fibril can be formed between the positively charged residue K28 and the COO^-^ at the C-terminal alanine A42. This salt bridge cannot be formed in DRI-Aβ_42_ peptides as the (negatively charged) C-terminal of the L-peptide becomes a (positively charged) N-terminus in the DRI-peptide, i.e., there will be a repulsive interaction between the NH3 groups on the N-terminal D-A42 and residue D-K28 (note that we count residues in DRI peptides starting from the C-terminus). While we saw in previous work that the K28-A42 salt-bridge is not crucial for the stability of S-shaped (L-)Aβ_42_ fibrils, its loss does lead to more flexibility of the β2-turn-β3 motif. ^20^ Hence, we conjecture that its replacement by a repulsive interaction in the DRI fibril lowers the stability below a critical threshold causing the beginning dissolution of the single-layer fibril as seen in our simulations.

In order to verify our assumption, we have constructed L and DRI versions of two Aβ_42_ mutants, and we have studied the stability of the corresponding single-layer fibrils in molecular dynamics simulations. We emphasize again that for simplicity we number residues in a DRI-peptide starting from the C-terminus instead, as usual in proteins, from the N-terminus. This allows us to keep the notation of the L-parent sequence. For example, the DRI analog of the K28E mutant is still called (DRI-) K28E, while with conventical counting it would become (DRI-)K15E. For the first mutant, K28A, we expect that L and DRI fibrils have similar stability, but one that is lower than seen for fibrils built from the L wild type. On the other hand, in the K28E mutant is a positively charged lysine replaced by a negatively charged glutamic acid, which should raise the stability of DRI fibrils to that of the L wild type, while lowering the stability of the L-fibril below that of the L wild type fibril. Representative configurations for this mutant are shown in Figure 6b and 6d.

**Table 3:** Relative fluctuation of the angle between β2 and β3 strands in single-layer fibrils made of L or DRI (wild type or mutant) Aβ_42_-peptides. Shown are averages and standard deviation (STD) of the angle and distance/number of contacts between residues 28 and 42.

| Single-layer | Number of sidechain contacts between residues 28-42 (inter-chain or intra-chain) | | Average distance between the sidechains of residues 28-42 / nm | | Angle between $\beta 2$ and $\beta 3$ strands/ degree | |
| --- | --- | --- | --- | --- | --- | --- |
|  | Average | STD | Average | STD | Average | STD |
| 7-L | 5.1 | 0.3 | 0.5 | 0.1 | 6.7 | 0.4 |
| 7-D | 1.9 | 0.3 | 1.7 | 0.3 | 72 | 19 |
| 7-L' | 2.2 | 0.2 | 1.4 | 0.2 | 49 | 15 |
| 7-D' | 3.7 | 0.5 | 0.9 | 0.1 | 24 | 6 |
| 7-L* | 1.3 | 0.2 | 1.5 | 0.2 | 45 | 13 |
| 7-D* | 4.9 | 0.3 | 0.5 | 0.1 | 6.5 | 0.6 |

Our results of both the wild type and the mutant simulations are listed in Table 2 and Table 3. In Table 2 we show the twist for the various cases. Note that the DRI-K28E mutant leads to a twist of similar magnitude than seen for the L wild type fibrils. In Table 3 we record the number of side chain contacts between residues 28 and 48, their average distance, and the angle between the β2 and β3 strands. The same quantities are shown for the double-layer model in supplemental Table S3. Here, a contact is defined by a minimal distance between heavy sidechain atoms that is less than 4.5 Å.

As expected, in the cases where residue 28 and the corresponding terminus have opposite charges, as in the case of the L-wild type and the DRI K28E mutant, the distance between the residue and the terminus is small and the number of contacts between the two large; i.e., the residue and the terminus are tightly connected by a salt bridge that is not formed when residue and terminus have the same charge or one of them is neutral. This salt bridge locks the β2 and β3 strands in place: the angle formed by the two strands stays below 10 degrees. On the other hand, while even without this salt bridge, the β2-strands keep their inter-chain hydrogen bonds, the angle between β2 and β3 fluctuates and can grow large, see Figure 6c and 6d. As a consequence, the β3 strands of the six chains in the fibril move against each other, reducing fibril stability and causing the observed large RMSD values in the wild type DRI fibril. We remark that our conjecture is also supported by simulations of fibrils with capped end groups (data not shown) where the single-layer L and DRI-fibrils have similar stability, sitting in between that of the stable non-capped L-fibrils and the unstable non-capped DRI fibrils.

#### 3.1.3 Incomplete symmetries cause disparities between L-peptide fibrils and DRI-peptide fibrils

Our above results have demonstrated that DRI-Aβ-peptides can assemble into similar fibrils than seen for the parent L-peptides, with only marginal differences in stability between L and DRI forms for double-layer Aβ_40_ and Aβ_42_-fibrils. Possible stability deviations are difficult to resolve as the double-layer architectures guarantee a large solidity that dwarfs the differences in interchain hydrogen bonding. Hence, only for the single-layer Aβ_42_-fibrils do we find a clear gap in stability between L-fibrils and DRI-fibrils. This gap results from interaction between residues close to the terminals and the charges of these terminals as a residue that is in the parent L-peptide close to the positively charged N-terminus will now be close to the negatively charged C-terminus, and vice versa. Hence, the lack of stability is due to a break in the symmetry between the L and DRI Aβ_42_-peptides, and full stability is recouped when this symmetry is recovered by the mutation K28E in the DRI-peptide. While not reducing fibril stability, opposite arrangement and orientation of the D23-K28 in the U-shaped Aβ_40_-chains, again breaking the symmetry between L and DRI fibril architecture, reduces the magnitude of the twist angle in double-layer DRI-Aβ_40_-fibrils. We have not investigated whether the effect would be more noticeable (and lead to a reduced stability) for single-layer DRI-Aβ_40_-fibrils as single-layer Aβ_40_-fibril structures have not been reported.

### 3.2 The effect of DRI Aβ-peptides on amyloid formation

When interacting with L-peptide oligomers and fibrils, the subtle differences in structure of DRI-Aβ peptides, resulting from interaction between charged residues close to the terminals and the charges of these terminals, from opposite arrangement and orientation of salt bridges, and from the different magnitude and orientation of twists of β-strands, may alter stability and propagation of amyloids. The possible outcomes could be either promotion or inhibition of fibril growth. In order to investigate the effect coming from presence of DRI-peptides in L-fibrils, we look into the stability of various hybrid fibril fragments made up out of mixtures of L and DRI peptides.

#### 3.2.1 Stability of hybrid double-layer Aβ_40_ fibrils

We start with the two-layer Aβ_40_ fibril where for the U-shaped chains the interaction between residues close to the termini and the terminal charges can be neglected. Hence, differences in hybrid fibril stability will likely result from the different twist preferences of β-strands. Besides the pure L and DRI fibrils, made of seven chains in each layer and denoted by us as (7-L/7-L) or (7-D/7-D), we also study hybrid fibrils where either one of the end chains is switched from a L-peptide to a DRI peptide (7-L/6-L:1-D), or where one in the middle is exchanged (7-L/3-L:1-D:3-L). The first case allows us to test variations in fibril growth by attachment of DRI peptides instead of L-peptides; while the second one probes the stability of a fibril that has a DRI peptide incorporated. An implicit assumption behind the choice of these two models is that the stability of pure fibrils (either DRI or L) is optimal and the incorporation or attachment of a DRI peptide in or to a L-fibril reduces stability. This assumption is by no means justified, and the opposite may be true, namely that U-shaped L and DRI Aβ_40_ chains like to attach to each other. In the latter case, fibrils formed by alternating L and DRI peptides would be most stable, and for this reason, we have also considered hybrids such as (L-D-L-D-L-D-L/D-L-D-L-D-L-D), for which we introduce the shorthand notation (2: L-D-L-D). All models are also listed in the supplemental Table S1.

With the same protocol as above, we have simulated these five systems with molecular dynamics, and the RMSD to the start configurations as function of time is shown in Figure 7(a). Not surprisingly, the differences are marginal as our work above already detected only small differences in stability between the L and DRI forms for the two-layer Aβ_40_ architecture with its U-shaped chains. Elongating a L-fibril by a DRI peptide (the 7-L/6-L:1-D system), the average number of hydrogen bonds is with 30(3) comparable to that of the L-fibril. However, the lone terminal DRI peptide in the 7-L/6-L:1-D model has on average 8 (2) hydrogen bonds while for the three terminal L-peptides the corresponding average is 12(2). Both values are comparable to that of the pure L and DRI fibrils, respectively. Hence, elongation of a L-fibril by a DRI peptide is energetically less favorable than elongation by a L-peptide by about five hydrogen bonds. On the other hand, there are on average about 28(4) hydrogen bonds connecting the DRI peptide with its L-neighbors in the 7-L/3-L:1-D:3-L model, higher than the corresponding number of 25(4) in a DRI fibril but lower than the number of 32(4) for an L-peptide in a L-fibril. Still, given the large fluctuations in the numbers, we conclude that once incorporated, a DRI-chain in an L-fibril has similar number of hydrogen bonds with its two neighbors than a L-peptide and the differences in stability between pure L-fibrils and hybrid fibrils become marginal. Within this picture the effect of DRI-peptides therefore would be to slow down elongation of fibrils but not or only marginally reducing their stability.

**Figure 7.**
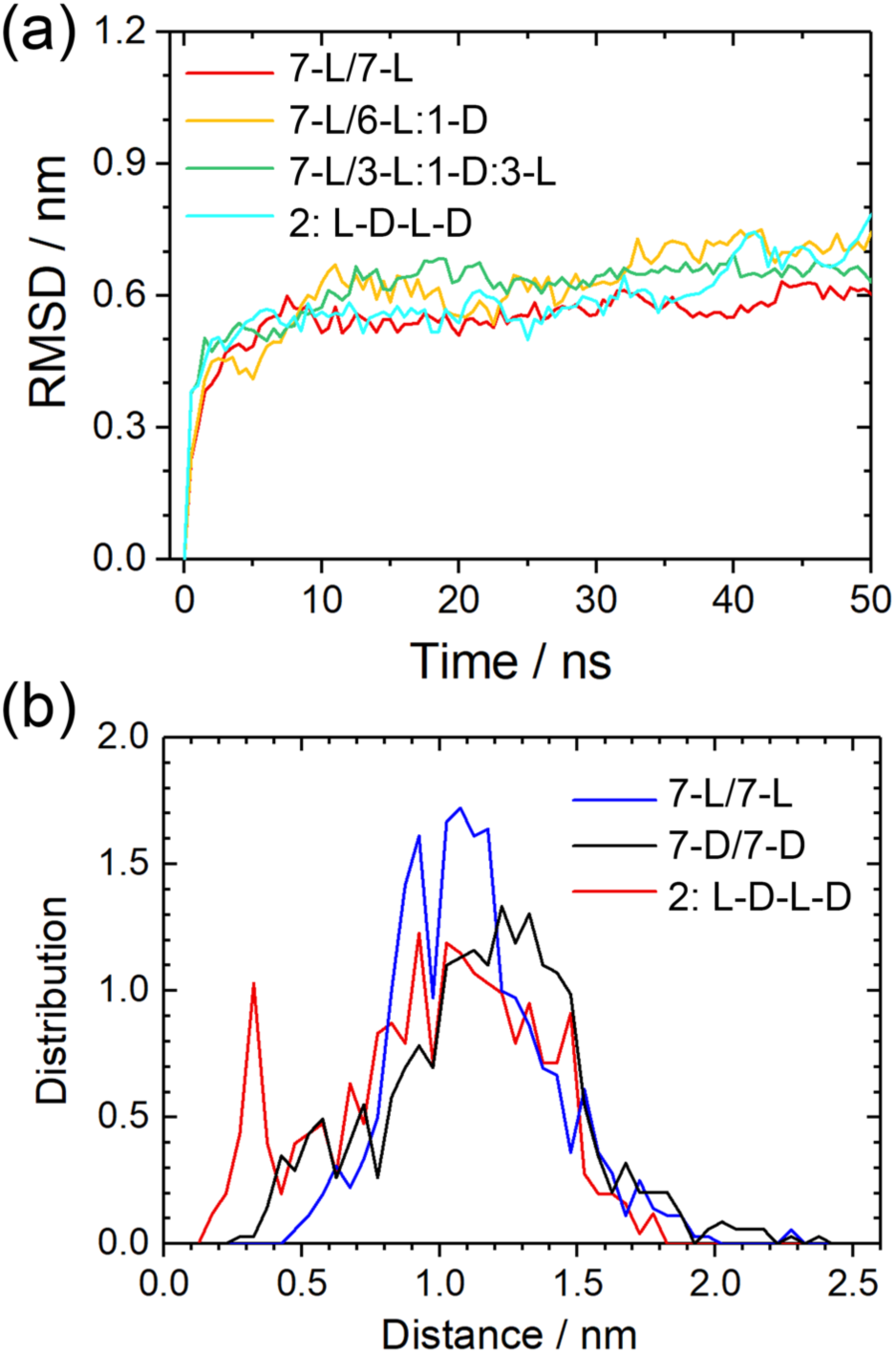
(a) Root-mean-square-deviation (RMSD) to the start configuration as function of time for all considered Aβ_40_ double-layer fibril models. Frequency of measured distances between atoms in adjacent terminals for double-layer Aβ_40_(b). For fibrils made solely from L-peptides, only the C-terminals are considered as the N-terminal region is disordered. Consequently, only the N-terminals are considered for DRI-fibrils, and for the hybrid L-D-L-D hybrid fibrils the C-terminals groups of L-peptide and N-terminal groups of DRI-peptides.

This scenario is supported by the hydrogen bond numbers seen for the 2:L-D-L-D model, where within the fibril each chain has on average 24(4) inter-chain hydrogen bonds with its neighbors, and at the fibril ends 9 (2) hydrogen bonds. These numbers demonstrate the difficulties that L and DRI-fibrils with this geometry have to attach to each other. The reasons can be understood from Figure 8. The location of hydrogen bonds for the parent L-fibril is shown in Figure 8a. This pattern is disrupted when a DRI chain is inserted, see Figure 8b, and new inter chain hydrogen bonds have to be formed that will result in a relative movement of the adjacent L and DRI peptides, as shown in Figure 7 (c, d). Snapshots from the various simulations show indeed that in all three Aβ_40_ hybrid models the DRI peptides shift slightly away from the turn region, with the β1 and β2 strands moving in opposite directions.

**Figure 8.**
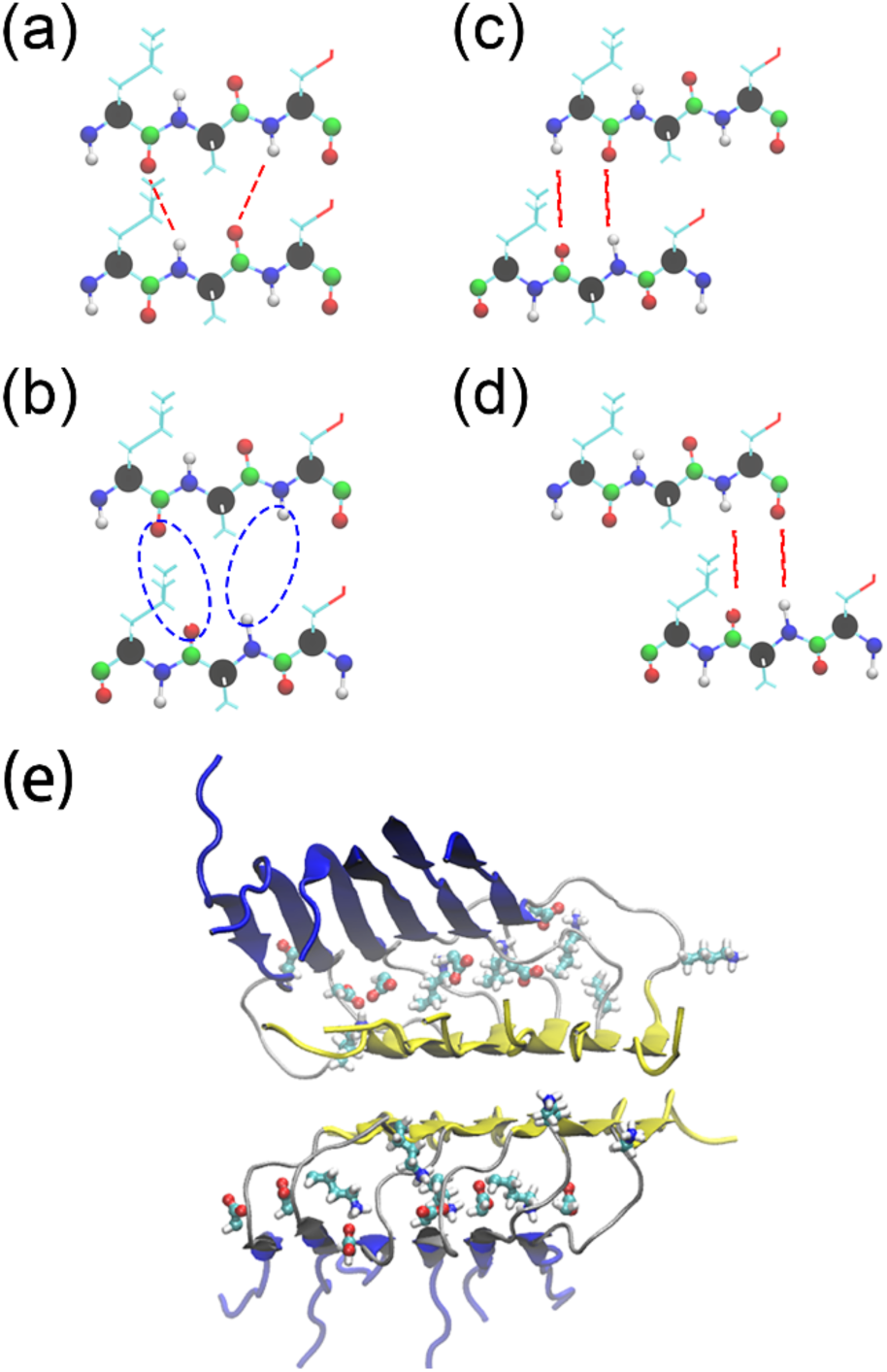
Interchain hydrogen-bonds in the L-Aβ_40_ fibrils. (a). This hydrogen bond arrangement is not stable when a L-peptide is replaced by a DRI peptide, see (b), and is replaced by either the one shown in (c) or the one shown in (d). The initial structure of a double-layer Aβ_40_ fibrils is presented in (e) with β1 marked in blue and β2 in yellow. The salt bridge forming side chains of residues D23 and K28 are drawn in an allatom representation.

#### 3.2.2 Stability of hybrid single-layer Aβ_42_ fibrils

The above problem may be less severe in Λβ_42_ fibrils, since the β2 and β3 strands are much shorter than the β1 strand. Hence, in order to see how the stability of hybrid fibrils depends on the geometry of the Aβ chains, we have looked also into the stability of Aβ_42_ fibrils. The individual chains of Aβ_42_ form S-shaped three-stranded configurations instead of the U-shaped configurations seen for Aβ_40_. As Aβ_42_ amyloids are more toxic than the more common Aβ_40_ amyloid_s_ it becomes important to know if added DRI-Aβ_42_ peptides have a different effect on promotion and inhibition of fibril growth than DRI-Aβ_40_ peptides have on growth of Aβ_40_ fibrils. In order to study this question, we have constructed also hybrid models Aβ_42_, and compared their stability with the pure L and DRI models.

Unlike Aβ_40_, Aβ_42_ peptides can assemble into single-layer fibrils, and hence, we study first the single-layer models 6-L:1-D; 3-L:1-D:3-L; and 1:L-D-L-D. Since, we have demonstrated earlier in this paper the role of the missing salt bridge between the K28 residue and the C-terminus in reducing the stability of fibrils made of DRI-Aβ_42_ peptides, and shown how the fibril stability is increased for the K28E mutant,^20^ we have considered two cases. In the first one, we use in the three hybrid models the wild type Aβ_42_ sequence for both the L-peptides and the DRI peptides, while in the second case we replace the wild type DRI-peptides by the K28E mutant DRI-peptide. In order to simplify our notation, we mark the mutant by the symbol D*. Hence, the additional three single-layer systems are 6-L:1-D*, 3-L:1-D*:3-L, and 1:L-D*-L-D*. All six hybrid fibril models are listed in supplemental Table S1.

The root-mean-square-deviation (RMSD) as function of time is for both sets of models shown in Figure 9, and representative figures are displayed in Figure 10. Note, that residues 1-10 are again not considered for calculating the RMSD as they form a flexible and disordered segment without defined structure. While the effect is smaller, we find as for the pure DRI models that RMSD values of the hybrid models are larger when the DRI peptides have the wild type sequence than when the DRI-peptide is a K28E mutant. Hence, as expected, the DRI-K28E mutant leads to a larger stability of the hybrid fibrils than when wild type DRI-Aβ_42_ are inserted, and for the 6-L:1D and 3-L:1-D:3-L models the stability is comparable to the pure L-fibrils. Similarly, comparable are the number of inter-chain hydrogen bonds which with 40-42 for the K28E mutant, and 39-41 for the wild type differ only little from the corresponding number of 42(3) for the L-fibril. However, the stability is significantly reduced for the 1:L-D-L-D model. where for the hybrid with mutant DRI peptides the final RMSD is about 0.8 nm compared the 0.6 nm seen for the pure L-fibril. This larger RMSD goes along with a reduced number of 38(3) interchain hydrogen bonds. The stability of the 1:L-D-L-D fibril is even lower when the DRI peptides have the wild type sequence, and correspondingly we find also a with 35(4) a smaller number of hydrogen bonds.

**Figure 9.**
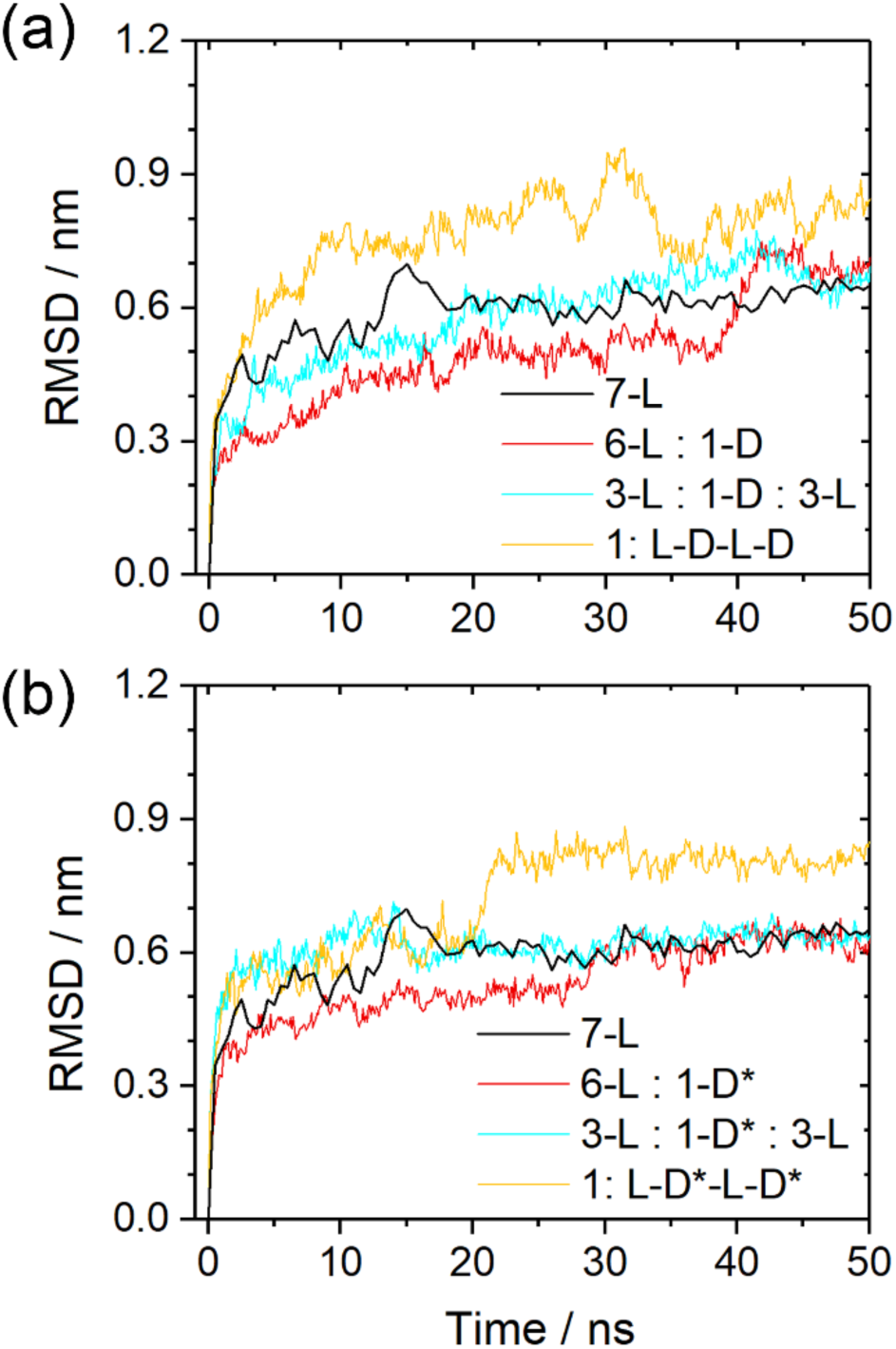
Root-mean-square-deviation (RMSD) to the start configuration as function of time for various hybrid single-layer Aβ_42_ models. In (a) we consider the case where both L-peptides and DRI-peptides have the wild type sequence, while in (b) only the L-peptides are wild-type Aβ_42_ while the DRI peptides are K28E mutants. For comparison, we show in both cases also the corresponding values for the pure wild type L-fibril.

**Figure 10.**
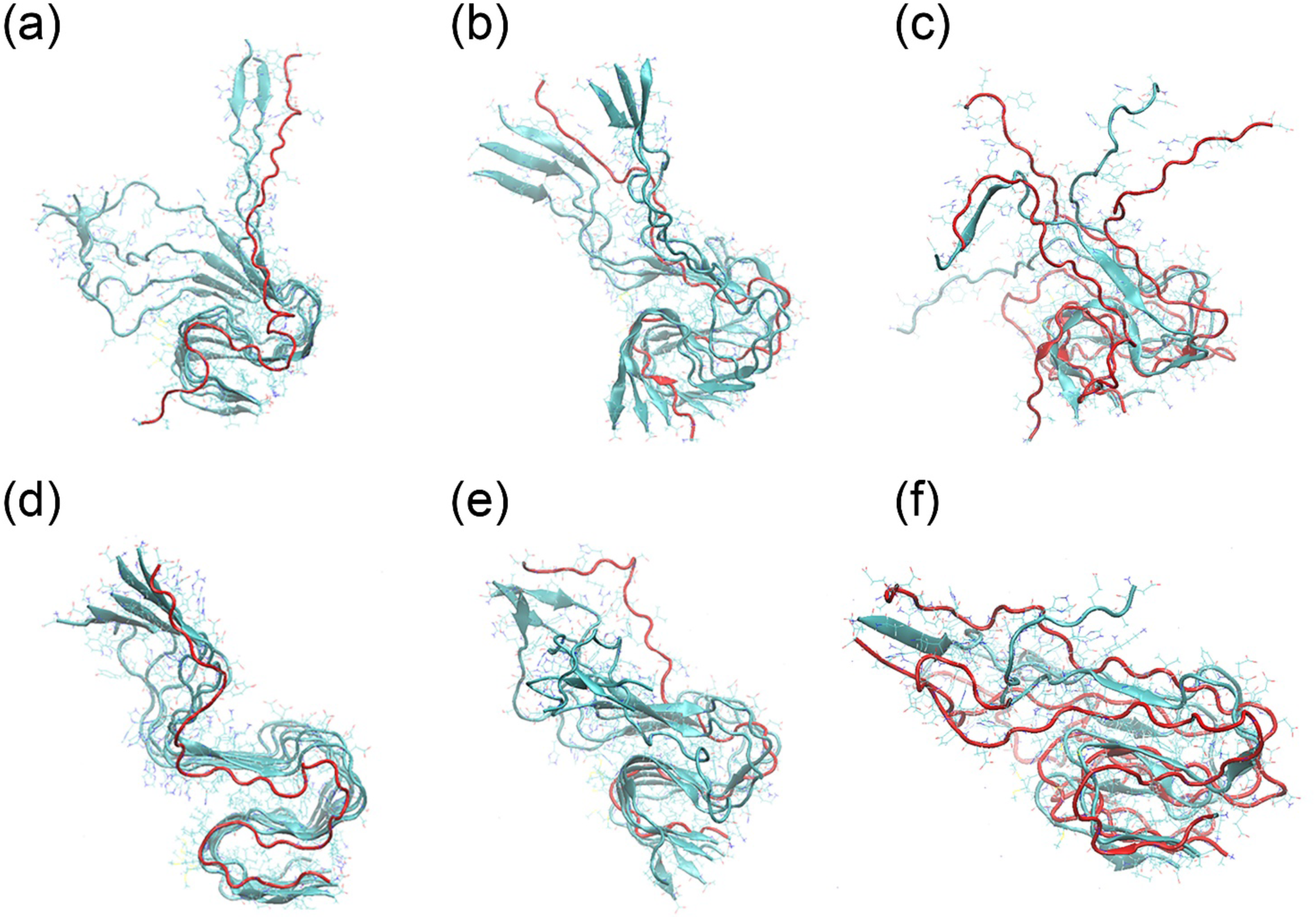
Representative conformations of various hybrid single-layer Aβ_42_ fibril models. In (a-c) both L and DRI peptides are wild type Aβ_42_ peptides (in (a) 6-L:1-D, (b) 3-L:1-D-3-L, and in (c) 1:L-D-L-D), while in (d-f) only the L-peptides have the wild type sequence but the DRI chains are K28E mutants (in (d) 6-L:1-D*, (e) 3-L:1-D*-3-L, and in (f) 1:L-D*-L-D*).

A similar picture is seen by the number of contacts between residues 28 and 42, their average distance and the angle between β2 and β3-strand, all shown in Table 4. As expected, the intra-chain salt bridge between residue 28 and the C-terminus is seen in all L-peptides, but it is formed in the DRI-peptides only for the K28E mutant. The stabilizing effect of the L-peptides reduces the sliding and the fluctuations of the angle between the β2 and β3-strand that lead to the large RMSD values seen for pure DRI-fibrils. This stabilizing effect is seen even in the case of wild type DRI-peptides in the hybrid fibrils. However, the stability of the hybrid fibrils stays below that of the pure L-fibrils.

As in the case of the Aβ_40_-fibrils with their U-shaped chains these results show that the presence of DRI-peptides has only little effect on a fully formed fibril, at least for the K28E mutant which similar to wild type L-Aβ_42_-peptides can form a salt bridge between residues 28 and the C-terminus. However, attachment of a DRI-peptide to a L-peptide located at the end of a fibril (or of a L-peptide to a DRI-peptide at the end of an existing fibril) appears to be energetically less favorable than elongation of L-fibrils by L-peptides.

**Table 4:**
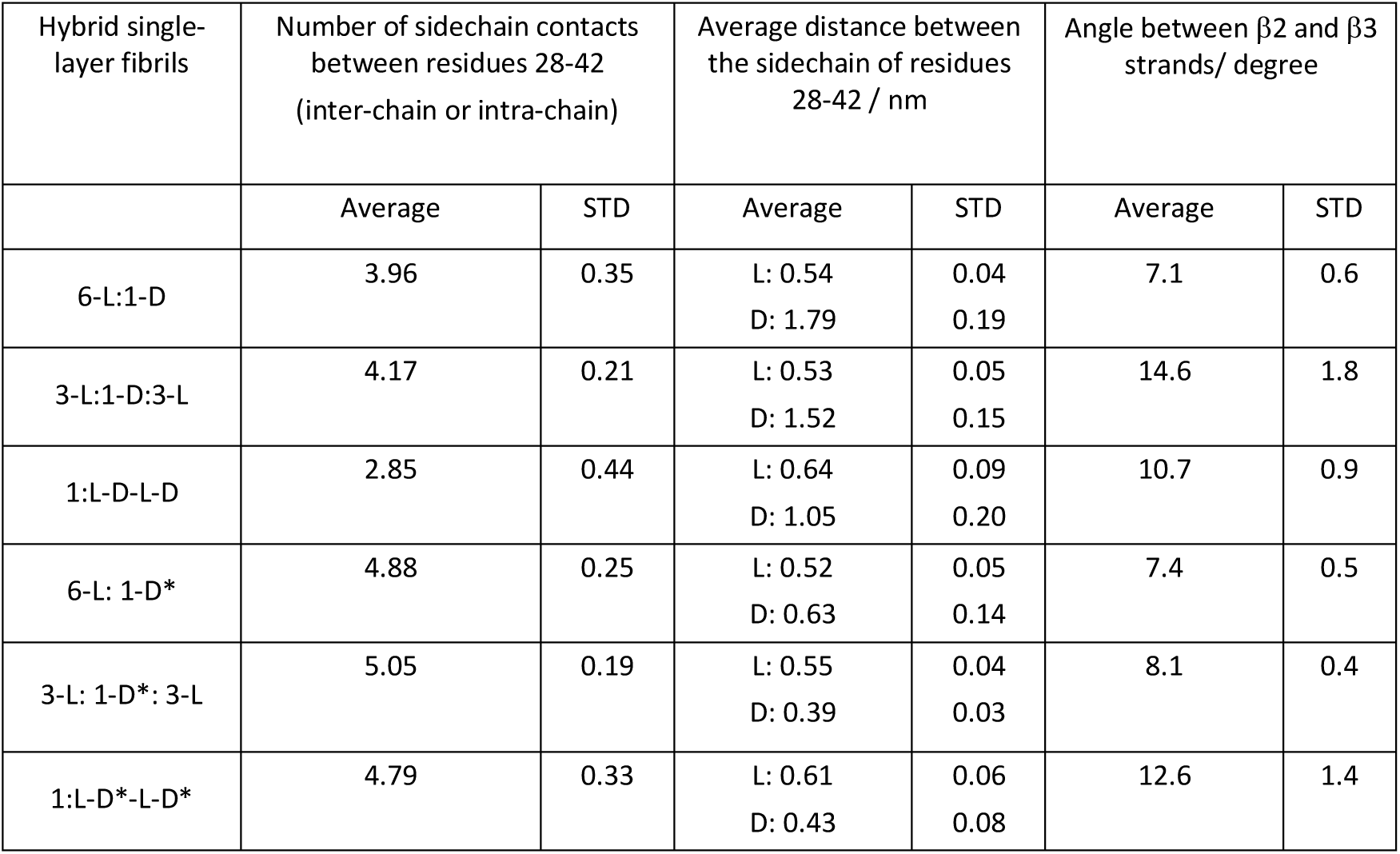
Relative fluctuation of the angle between β2 and β3 strands in hybrid single-layer fibrils made of L and DRI (wild type or mutant) Aβ_42_-peptides. Shown are averages and standard deviation (STD) of the angle and distance/number of contacts between residues 28 and 42.

| Hybrid single-layer fibrils | Number of sidechain contacts between residues 28-42 (inter-chain or intra-chain) | | Average distance between the sidechain of residues 28-42 / nm | | Angle between $\beta 2$ and $\beta 3$ strands/ degree | |
| --- | --- | --- | --- | --- | --- | --- |
|  | Average | STD | Average | STD | Average | STD |
| 6-L:1-D | 3.96 | 0.35 | L: 0.54<br>D: 1.79 | 0.04<br>0.19 | 7.1 | 0.6 |
| 3-L:1-D:3-L | 4.17 | 0.21 | L: 0.53<br>D: 1.52 | 0.05<br>0.15 | 14.6 | 1.8 |
| 1:L-D-L-D | 2.85 | 0.44 | L: 0.64<br>D: 1.05 | 0.09<br>0.20 | 10.7 | 0.9 |
| 6-L: 1-D* | 4.88 | 0.25 | L: 0.52<br>D: 0.63 | 0.05<br>0.14 | 7.4 | 0.5 |
| 3-L: 1-D*: 3-L | 5.05 | 0.19 | L: 0.55<br>D: 0.39 | 0.04<br>0.03 | 8.1 | 0.4 |
| 1:L-D*-L-D* | 4.79 | 0.33 | L: 0.61<br>D: 0.43 | 0.06<br>0.08 | 12.6 | 1.4 |

#### 3.2.3 Stability of hybrid double-layer Aβ_42_ fibrils

The magnitude of the effect is difficult to compare with the one observed for the Aβ_40_ fibrils, as the later are two-layered, with the interaction between the two-layer providing an extra stabilizing force. For this reason, we have in a second step extended our study of hybrid Aβ_42_-fibrils with S-shaped chains to double-layer assemblies. Because the single-layer investigations confirmed the stabilizing effect of choosing as the DRI-peptides the K28E mutant instead of the Aβ_42_ wild type, we have considered here only the three models 7-L/6-L:1-D*, 7-L/3-L:1-D*:3-L and 2:L-D*-L-D*. The root-mean-square-deviation (RMSD) as function of time is for both sets of models plotted in Figure 11, where also representative figures are displayed as Figure 11 (c)-10(e).

These configurations and plots indicate that the stability of all hybrid fibrils with mutant sequence for the DRI-peptides is comparable with that of the parent L-fibril. The number of interchain hydrogen bonds also does not differ, expect in the case of 2:L-D*-L-D*, where only 40 (4) instead of 43 (3) hydrogen bonds per chain are observed. Note that the differences in RMSD and hydrogen bonding are much smaller than for the single-layer fibril with alternating L and DRI peptides. This is because the alternation of L-peptides and K28E-DRI peptides leads to two-layer fibrils with a packing surface (52.3 nm^2^) between the layers that is about 19% larger than the packing surface (43.8 nm^2^) in pure L-Aβ_42_-fibrils, while the overall solvent accessible surface area increases only by 11% from a value of 194 nm^2^ (pure L-Aβ_42_-fibrils) to 215 nm^2^ (L-D*-L-D*). The tighter interaction between the two layers stabilize the fibrils and compensate for the loss of hydrogen bonding. The overall effect seems to be that presence of K28E-DRI-peptides does not reduce the stability of (wild type) two-layer L-Aβ_42_-fibrils. Figure 11 a seems to indicate that stability is even slightly increased. Similarly, neither is attachment of a K28E DRI-peptide to a L-fibril disfavored, nor if it happens, does it slow down further elongation, as alternating patterns are not discouraged by unfavorable binding.

**Figure 11:**
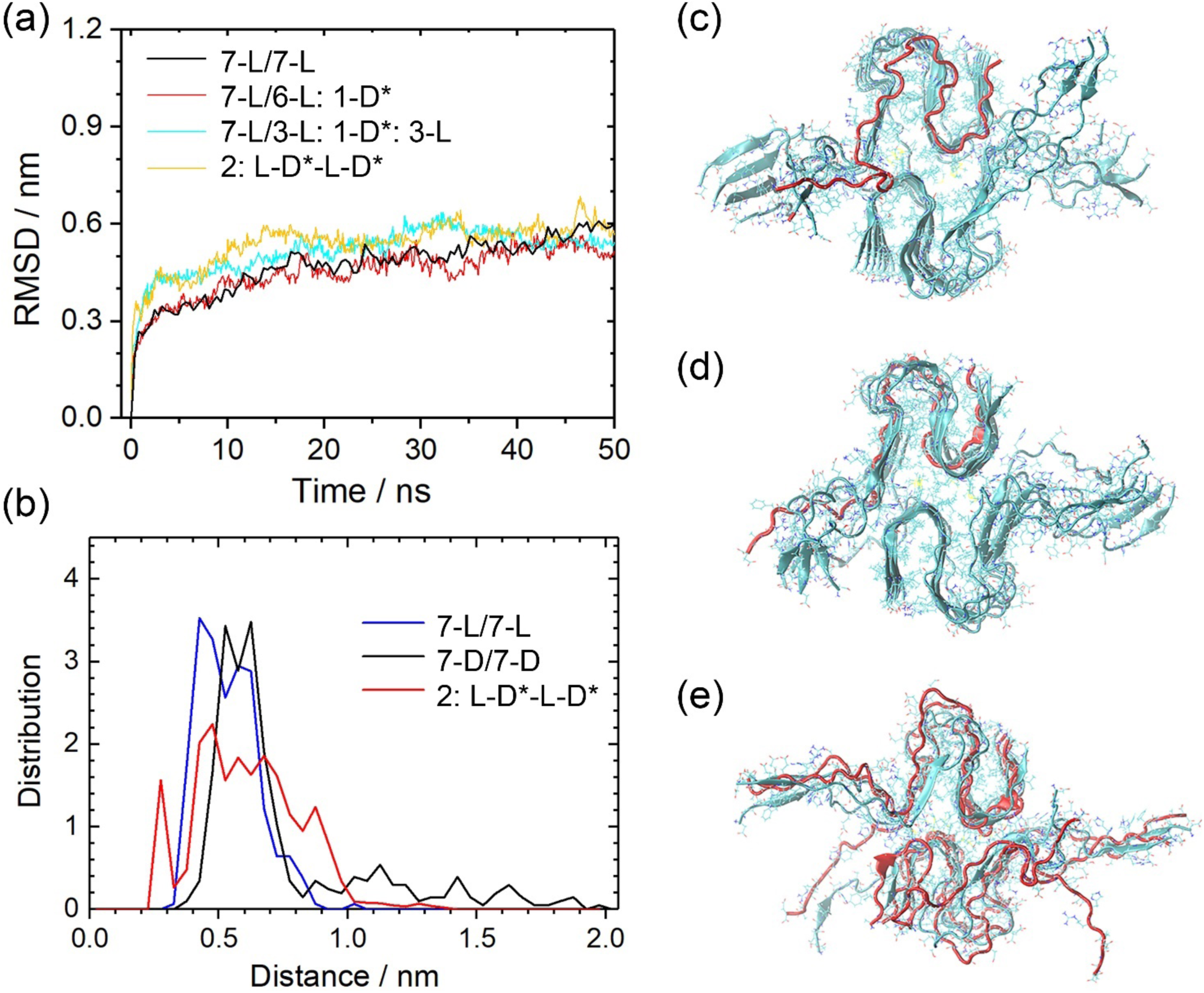
Root-mean-square-deviation (RMSD) to the start configuration as function of time. (a) and frequency of measured distances between atoms in adjacent terminals (b) for various hybrid double-layer Aβ_42_ models. In these fibril models have the L-peptides the wild type sequence while the DRI peptides are the K28E mutant. We show representative configurations for the fibril models 7-L/6-L:1D* (c), 7-L/3-L:1-D*:3-L (d) and 2:L-D*-L-D* (e).

It is surprising that the stabilizing effect of alternating pattern of L and DRI peptides is not more pronounced as the lower number of inter-chain hydrogen bonds should be compensated by two other effects. The first one is that L-peptides and DRI-peptides form anti-parallel β-sheets, while in pure L-fibrils or pure DRI-fibrils the chains form parallel β-sheets. The consequence is a tighter and more energetically favorable packing of side chains, see Figure 12a) and b), and a distribution of hydrogen bond distances that is shifted towards smaller values (and therefore stronger bonds), see Figure 12c.

Another effect of the anti-parallel arrangement of the L and DRI chains are attractive electrostatic interactions between the positive charge on the N-terminal of a DRI peptide with the negative charge on the C-terminal of a neighboring L-peptide, forming an additional fibril-stabilizing salt-bridge. The existence of such salt bridge can be seen in Figure 7(b). The distributions of the L and DRI fibrils differ little and are both centered around an average of 1.2 nm. On the other hand, a second peak (representing 5% of the population) is seen for the 2:L-D*-L-D* fibrils at 0.35nm - 0.4 nm, as a typical salt-bridge distance. Note that formation of a second possible inter-chain salt bridge, between the N-terminus of the L-peptide and the C-terminus of the DRI-peptide is less likely as the first (last) 10 residues in the L(DRI-)peptide are flexible and do not adopt a defined structure.

Both effects are more prominent in hybrid Aβ_42_-fibrils than in Aβ_40_-hybrids where they are counteracted by the opposite arrangement and orientation of the characteristic salt bridge D23-K28 salt bridge and the different magnitude and orientation of β-strands twists. In hybrid Aβ_42_-fibrils, on the other hand, the weaker hydrogen bonding could be compensated (or even exceeded) by the more favorable anti-parallel packing and the additional salt bridge. This salt bridge would lock-in the terminal regions, stabilizing in this way the fibril, and guides the attachment of new chains speeding up fibril elongation. While our data confirm the presence of both of these additional stabilizing effects, their magnitude is not large enough to compensate for the less favorable hydrogen bonding between L and DRI peptides in single-layer hybrid Aβ_42_-fibrils. It is, however, visible in the case of double-layer hybrid Aβ_42_-fibrils where the loss of hydrogen bonding is less severe. Presence of DRI-peptides appears even to stabilize the fibrils, potentially shifting the equilibrium away from the more toxic oligomers to less-toxic fibrils. Note that both effects may also lead to faster attachment of chains, and a faster formation of seeds needed to start nucleation of fibrils. Our present stability investigations do not allow us to study these kinetic effects but in future work we plan to study the differences in dimerization of two L-Aβ (or two DRI-Aβ) peptides with that of a L-Aβ and a DRI-Aβ peptide. If our conjecture is correct we would expect a much faster dimerization rate for the mixed system than for the pure ones.

**Figure 12.**
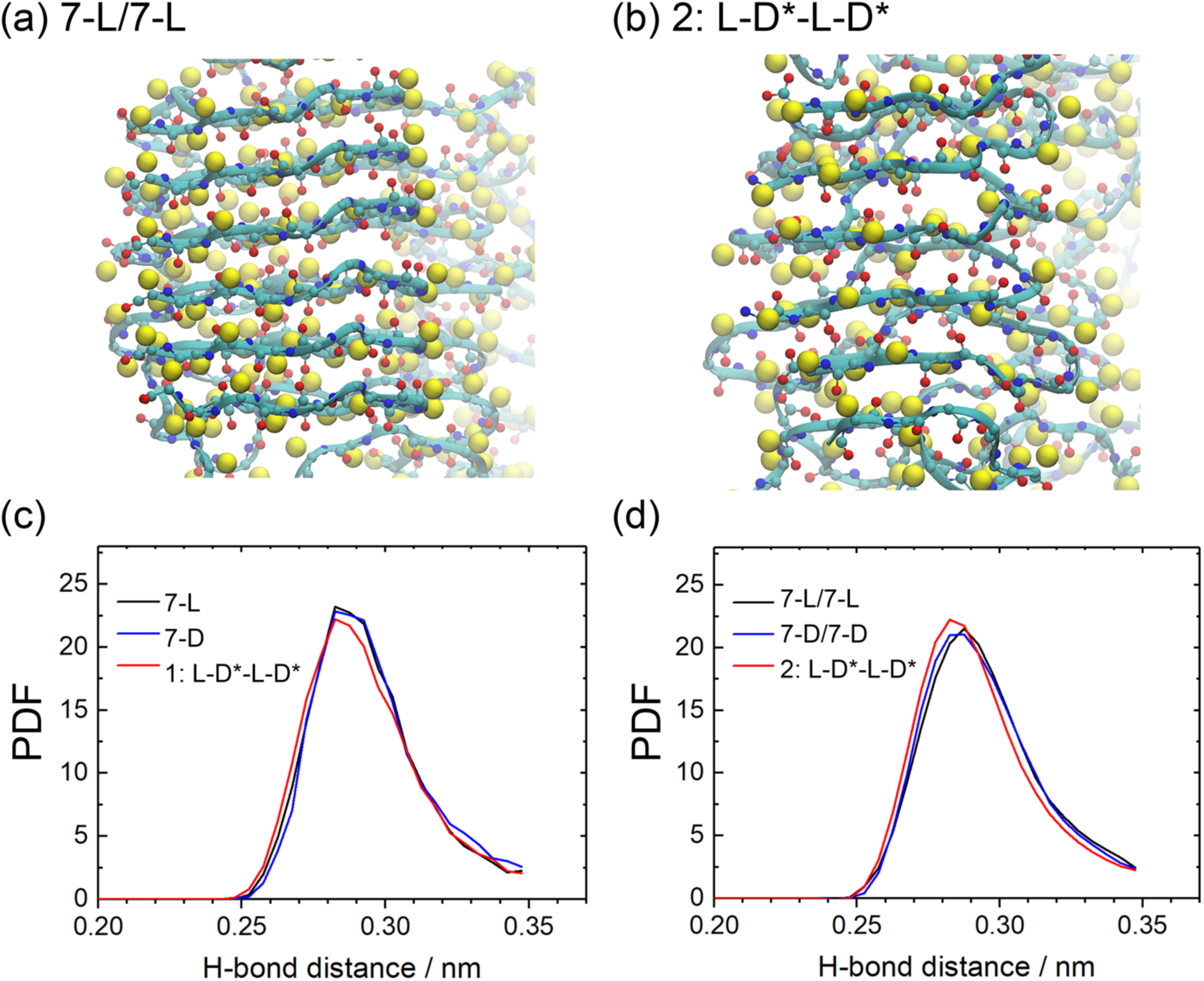
The backbone of 7-L/7-L (a) and 2:L-D-L-D (b) Aβ_42_ models. Of the sidechains only the Cβ atom are shown, drawn as yellow balls. The 7-L/7-L fibrils have a typical parallel β-sheet arrangement with the sidechains aligned, while they are alternating in the anti-parallel β-sheet pattern of the 2:L-D*-L-D* fibril. The distribution (PDF) of mainchain hydrogen-bond length for single-layer(c) and double-layer Aβ_42_(d).

## 4. Conclusions

Using molecular dynamics simulations, we have studied how full-length D-retro-inverso Aβ peptides (DRI-Aβ_40_ and DRI-Aβ_42_) interfere with fibril formation. Our simulations rely on a scheme to generate DRI-peptide configurations from the L-parent that is described in detail in the method section. The potential use of these peptides as drug candidates is motivated by the assumption that they can form stable hybrid aggregates with L-Aβ-peptides. This is likely as the mirroring of the sequence together with the replacement of L-amino acids by D amino acids leads to molecules that resemble in structure and stability their L-parent.

However, there are subtle differences that result from incomplete symmetries between L-fibrils and DRI-fibrils, i.e., mirroring the sequence of amino acids in Aβ peptides and replacing L amino acids by their D enantiomers does not preserve exactly the parent structure. For this reason, we have first studied the stability of typical Aβ-fibril models where the L-Aβ peptides are replaced by the corresponding DRI-Aβ peptides. We find that neighboring chains in DRI-Aβ_40_ fibrils share less hydrogen bonds than they do in the L-fibrils, leading to reduced stability of the DRI fibrils. They also have a twist of β-sheets that varies not only in sign but also in magnitude from the ones seen in the L-assemblies. We explain these differences with the change in the staggering of the salt bridge D23-K28, characteristic for the U-shaped Aβ_40_, from (i,i+2) in L-peptides to (i, i-2) in DRI peptides. Presence of this salt bridge restricts the formation of hydrogen bonds involving residues 23 to 29 between chains (i,i+1) in the DRI fibril, but because of the different staggering not in L-fibrils. This effect is not seen for the Aβ_42_ fibril with their S-shaped chains where this salt bridge does not exist. However, in wild type DRI-Aβ_42_ fibril the salt bridge, seen in L-fibrils between K28 (carrying a positive charge) and the negatively charged C-terminus, does not exist. Instead it is replaced by a repulsive interaction by the corresponding (D-) residue with the positively charged N-terminus, which, especially in the single layer fibril, lowers again the stability of the fibril. Consequently, the mutation K28E in the DRI-Aβ_42_ peptides lead to fibrils whose stability is comparable with the L-parents.

We then have studied the effect of DRI-Aβ peptides on the stability of fibrils simulating various hybrid assemblies of DRI-Aβ peptides with L-peptides. We find that the stability of hybrid Aβ_40_ fibrils is visibly smaller than the one of the pure L-fibrils. The differences are small, but the adverse effect may be suppressed by the large intrinsic stability of the double-layer arrangement. On the other hand, even for the single layer Aβ_42_ fibrils are the stability differences between pure L-fibrils and hybrid fibril marginal as long as the DRI peptide is the mutant K28E. Surprisingly does the alternating pattern of L and DRI peptides in the 1:L-D*-L-D* hybrid does not stabilize the fibril but even reduces its stability. Hence, the anti-parallel arrangement of the L and DRI chains, leading to an attractive electrostatic interaction between the positive charge on the N-terminal of a DRI peptide with the negative charge on the C-terminal of a neighboring L-peptide, cannot compensate for the loss of hydrogen bonding in the hybrid fibril. This is different for the double-layer Aβ_42_-fibrils where the alternating pattern even increases the contact between the two layers. While this effect is small, the hybrid fibrils have here even a higher stability than the L-fibrils. We speculate that cross-seeding of L and DRI-peptides may even ease the kinetics of fibril formation encouraging faster attachment of chains, and a faster formation of seeds needed to start nucleation of fibrils.

Given the only slightly larger stability of hybrid fibrils, we conjecture that cross fibrilization of Aβ_42_ L‐ and DRI-peptides is likely, and it may be even kinetically favorable. This implies that these full-length DRI-peptides may enhance the fibril formation and decrease the ratio of soluble toxic Aβ oligomers. An important caveat is that one has to choose the equivalent to the K28E mutant as DRI-peptides as otherwise an important salt bridge cannot be formed. The situation is less clear for the case of Aβ_40_ where the stability of the hybrid fibrils is below that of pure L-fibrils. Cross-seeding appears here less likely, but as the stability differences are marginal, still possible. Hence, our data suggest that full-length DRI-peptides may enhance Aβ-fibril formation and decrease the ratio of soluble toxic Aβ oligomers, pointing out a potential for D-amino-acid-based drug design targeting Alzheimer's disease.

## Acknowledgements

Simulations were done on the SCHOONER cluster of the University of Oklahoma and the Extreme Science and Engineering Discovery Environment (XSEDE) which is supported by the National Science Foundation (NSF). Computing time was awarded under grant MCB160005. We acknowledge financial support from National Institutes of Health (NIH) under research grant GM120634. We thank Jun (Eva) Yi for suggesting this project, initial discussions, and reading the final manuscript.

